# CellMeSH: Probabilistic Cell-Type Identification Using Indexed Literature

**DOI:** 10.1101/2020.05.29.124743

**Authors:** Shunfu Mao, Yue Zhang, Georg Seelig, Sreeram Kannan

## Abstract

Single-cell RNA sequencing (scRNA-seq) is widely used for analyzing gene expression in multi-cellular systems and provides unprecedented access to cellular heterogeneity. scRNA-seq experiments aim to identify and quantify all cell types present in a sample. Measured single-cell transcriptomes are grouped by similarity and the resulting clusters are mapped to cell types based on cluster-specific gene expression patterns. While the process of generating clusters has become largely automated, annotation remains a laborious ad-hoc effort that requires expert biological knowledge. Here, we introduce CellMeSH - a new automated approach to identifying cell types based on prior literature. CellMeSH combines a database of gene-cell type associations with a probabilistic method for database querying. The database is constructed by automatically linking gene and cell type information from millions of publications using existing indexed literature resources. Compared to manually constructed databases, CellMeSH is more comprehensive and scales automatically. The probabilistic query method enables reliable information retrieval even though the gene-cell type associations extracted from the literature are necessarily noisy. CellMeSH achieves up to 60% top-1 accuracy and 90% top-3 accuracy in annotating the cell types on a human dataset, and up to 58.8% top-1 accuracy and 88.2% top-3 accuracy on three mouse datasets, which is consistently better than existing approaches.

**Availability:** Web server: https://uncurl.cs.washington.edu/db_query and API: https://github.com/shunfumao/cellmesh

## Introduction

Single-cell RNA sequencing (scRNA-seq) is providing an unprecedented resolution in understanding cellular heterogeneity at the single-cell level, and offering novel biological insights into multi-cellular organisms [1–18]. A key step required in order to enable the aforementioned applications is cell-type identification that annotates cells with biologically meaningful cell types. However, cell-type annotation remains primarily a manual process and automatic cell-type identification is an important open problem [19].

One line of automatic cell-type identification methods [20–25] (Table 1) annotates *clusters of cells* obtained via a standard scRNA-seq workflow. However, the range of cell types that can be annotated as well as the accuracy of these methods remain insufficient. A typical scRNA-seq analysis workflow [26–29] begins with the preparation of gene-cell expression matrix (a gene-cell expression matrix is obtained from the rawreads after a sequence of steps such as read quality control [30], alignment [31, 32] and quantification [33, 34]). This matrix is used as a starting point on which clustering [35–39], dimension reduction [40, 41], and differential expression analysis [42] are further applied giving rise to a set of genes that are expressed specific to a cluster (which we refer to as, cluster differentially expressed genes). To annotate clusters with cell types, these existing methods [20–25] use the cluster differentially expressed genes to query databases, that connect genes to cell types. The databases are collected either from a few specific studies [20, 21], from manual literature surveys [22–24], or from scRNA-seq experiments which have their clustered cells pre-annotated according to the cell-type markers manually compiled from literature [25]. The database query mechanisms can return a list of unsorted cell types [21, 22, 25] or a list of cell types sorted by their statistical significance with the query genes, essentially based on Fisher’s exact test [20, 23] or a Kolmogorov-Smirnov test [24]. The common issue for these cell-type identification methods is that their databases are not comprehensive; more critically it is also laborious to update and expand them.

**Table 1.** Existing Works for Cell-type Identification. They can differ in multiple aspects: the Type (if the work provides a webserver, a general method, or a user-side software), the Target (if the work can be used to annotate cell types for clusters or for cells directly), the Query Input (if a gene set or gene expressions will be used to represent the query cell), the Query Method (if the results are retrieved by enumeration, statistical tests, or machine learning inferences), the Database (from what sources the gene-cell knowledge come from) and the Query Output (how the retrieved cell types can be provided).

| Name | Type | Target | Query Input | Query Method | Database | Query Output |
| --- | --- | --- | --- | --- | --- | --- |
| CellMeSH | webserver | cluster | gene set | maximum likelihood estimation | indexed literature (MEDLINE [49] MeSH [51] cells and Gene2pubmed [50] genes) | ranked list of cell types |
| CellFinder (2013) [21] | webserver | cluster | gene set | enumeration | mainly from existing ontologies, augmented by text mining of literature with manual verification | list of cell types |
| CellMarker (2018) [22] | webserver | cluster | gene set (actually a single gene) | enumeration | manual literature survey | list of pairs of tissue and cell type |
| PanglaoDB (2019) [25] | webserver | cluster | gene set | enumeration | scRNA-seq experiments (cell clusters were pre-annotated according to hand-curated cell-type markers) | list of labeled cell clusters, and a barplot for frequencies of the labels |
| Enrichr (2013) [20] | webserver | cluster | gene set | modified Fisher exact test | Mouse and Human Gene Atlases [59], CCLE [67] and NCI-60 [68] | ranked list of cell types |
| Hypergeometric test[23] | method | cluster | gene set | Fisher's exact test | need manual compile of cell and marker genes | ranked list of cell types based on raw enrichment scores |
| GSVA (2013) [24] | method | cluster | gene expression | Kolmogorov Smirnov random walk statistic | need manual compile of cell and marker genes | ranked list of cell types based on raw enrichment scores |
| scQuery (2018) [43] | webserver | cell (or optionally cluster) | gene expression (or optionally gene set) | nearest neighbor of neural embedding | scRNA-seq experiments from GEO and ArrayExpress | ranked list of cell types |
| scMatch (2019) [44] | software (Python) | cell | gene expression | correlation (Spearman or Pearson) | annotated gene expression data | ranked list of cell types based on raw scores |
| ACTINN (2019) [45] | software (Python) | cell | gene expression | neural network | annotated gene expression data | ranked list of cell types based on raw scores |
| SingleCellNet (2019) [46] | software (R) | cell | gene expression | random forest classification [69] | annotated gene expression data | labeled single cells |
| Garnett (2019) [47] | software (R) | cell | gene expression | supervised classification | need manual compile of cell and marker genes | labeled single cells |
| cellassign (2019) [48] | software (R) | cell | gene expression (critical to use only marker genes) | maximum a posteriori estimation using EM algorithm [70] | cell and marker genes from bulk RNA-seq experiments or literature | ranked list of cell types |

Another line of recent work [43–48] (Table 1) predicts cell *types for single-cells* (rather than clusters) using the gene-cell expression matrix directly. However, these methods require either existing annotated gene expression profiles [43–46] or hand-curated celltype marker-gene files [47, 48] as prior knowledge. The majority of these methods follow a machine learning approach, by first training a model based on the prior knowledge, and then utilizing the trained model either to classify the input gene expression vector to a reference cell type [45–47], or to project the input gene expression vector to an embedding vector and match to the reference cell type that has the most similar embedding [43]. Some of these methods follow a more statistical approach, by annotating the input gene expression vector with the reference cell type that has the highest correlation [44] or maximum a posteriori estimation score [48] for the input. All of these methods are unable to annotate the cell types that are either unseen in prior experiments (as used in [43–46]) or absent from the marker file which typically contains only a small number of known cell types (as used in [47, 48]).

The main goal of this paper is to address the shortcomings of the existing cell-type identification methods (Table 1) by exploiting the existing indexed literature resources such as MEDLINE [49] and Gene2pubmed [50]. We particularly focus on celltype annotation at the cluster level [20–25]. MEDLINE and Gene2pubmed respectively specify important Medical Subject Heading (MeSH) [51] concepts (a set of hierarchically-organized biological terms, including cell types) and NCBI genes [50] for a large class of biomedical publications. A natural approach is then to build a database that connects these genes with MeSH cell types. Since the genes and cell types are indexed for a large class of publications, the database forms a rich resource in associating genes with cell types. Furthermore, since the underlying resources (MEDLINE and Gene2pubmed) expand as new papers come up, the extracted database can also be automatically updated. However, connecting these genes and MeSH cell types simply based on the number of papers where they co-occur results in (a) many spurious gene-cell relationships, and (b) biases due to the widely varying number of publications mentioning a gene or cell-type. Existing query methods [23, 24] may not work well for such a noisy database, because they all implicitly assume that the database is noiseless and has only true gene-cell associations. Therefore, utilizing these literature resources necessitates the design of novel query methods.

Here, we propose CellMeSH (Cell-type annotation with MeSH terms), a new method to annotate clustered single-cell data, comprising of two key parts: a database of gene-cell type mappings, and a novel query method. Its accompanying web server and open-source API are able to take an input of a set of genes (such as the differentially expressed genes of a cluster of cells), and output a list of candidate cell types sorted by their relevance to the genes (Fig. 1). Unlike many of the methods that assign cell types to cells using gene cell expressions directly, CellMeSH does not need a separate training dataset.

There are two key innovations in CellMeSH. First, CellMeSH builds its database in a scalable way, by automatically linking the genes (as indexed in Gene2pubmed) and MeSH cell types (as indexed in MEDLINE) from millions of publications. Such large-scale gene-cell linking makes the database more comprehensive and easy to expand when new literature comes online. Second, to address the challenges of publication bias and potentially error-prone gene-cell associations in building the database, we develop a novel probabilistic database query method using maximum likelihood estimation.

**Figure 1.**
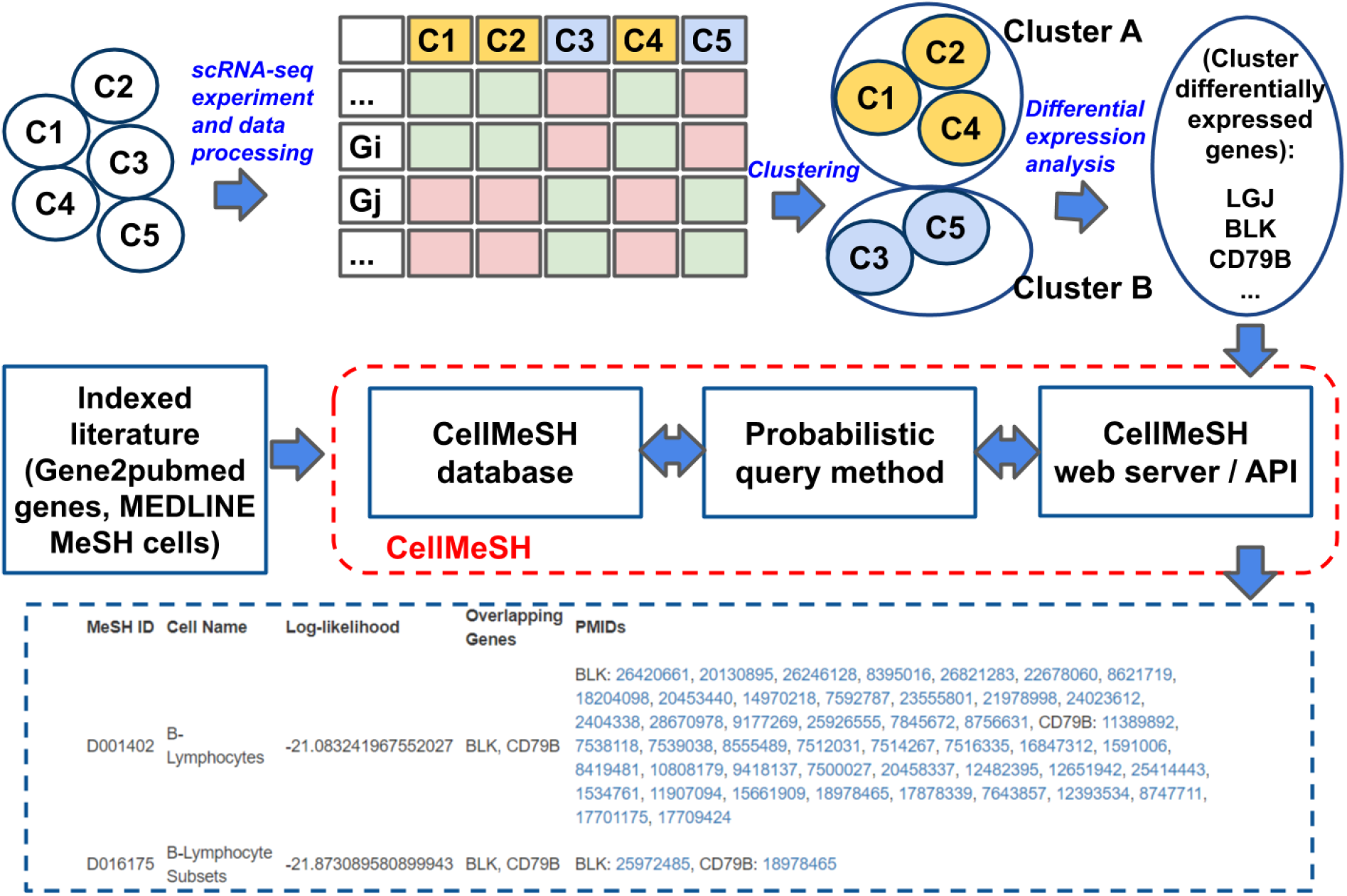
CellMeSH Overview. CellMeSH mainly addresses the annotation of cell types, which is usually the last step of scRNA-seq analysis. It provides web server and API to take input of the differentially expressed genes of clustered cells, and produce output including ranked candidate cell types, overlapping genes and related literature resources. It relies on a novel database by linking the Gene2pubmed genes and MEDLINE MeSH cell types. The database is queried by a novel probabilistic method based on maximum likelihood estimation.

Through a variety of experiments on human and mouse scRNA-seq datasets, we demonstrate that CellMeSH has richer information in its database linking genes and cell types, a robust query method, and an overall better annotation performance than existing methods.

Below, we first go through the key parts of CellMeSH including the database and query method. We next demonstrate the superior annotation performance of CellMeSH for human and mouse scRNA-seq datasets. We then describe the CellMeSH web server and its open-source API. Lastly, we discuss our future work.

## Results

### CellMeSH database

To construct the CellMeSH database, we first filter MEDLINE for references containing MeSH cell types (Fig. 2). MEDLINE [49] is a bibliographic database containing around 30 million references to biomedical and life science journal articles, including to most articles in PubMed. Each MEDLINE reference associates a subset of terms from Medical Subject Headings (MeSH) [51] with each article. MeSH includes 570 terms related to cell types (nested under the MeSH category “Cell” with tree number A11). Filtering MEDLINE for MeSH cell types results in a reduced dataset of 3.8M articles.

**Figure 2.**
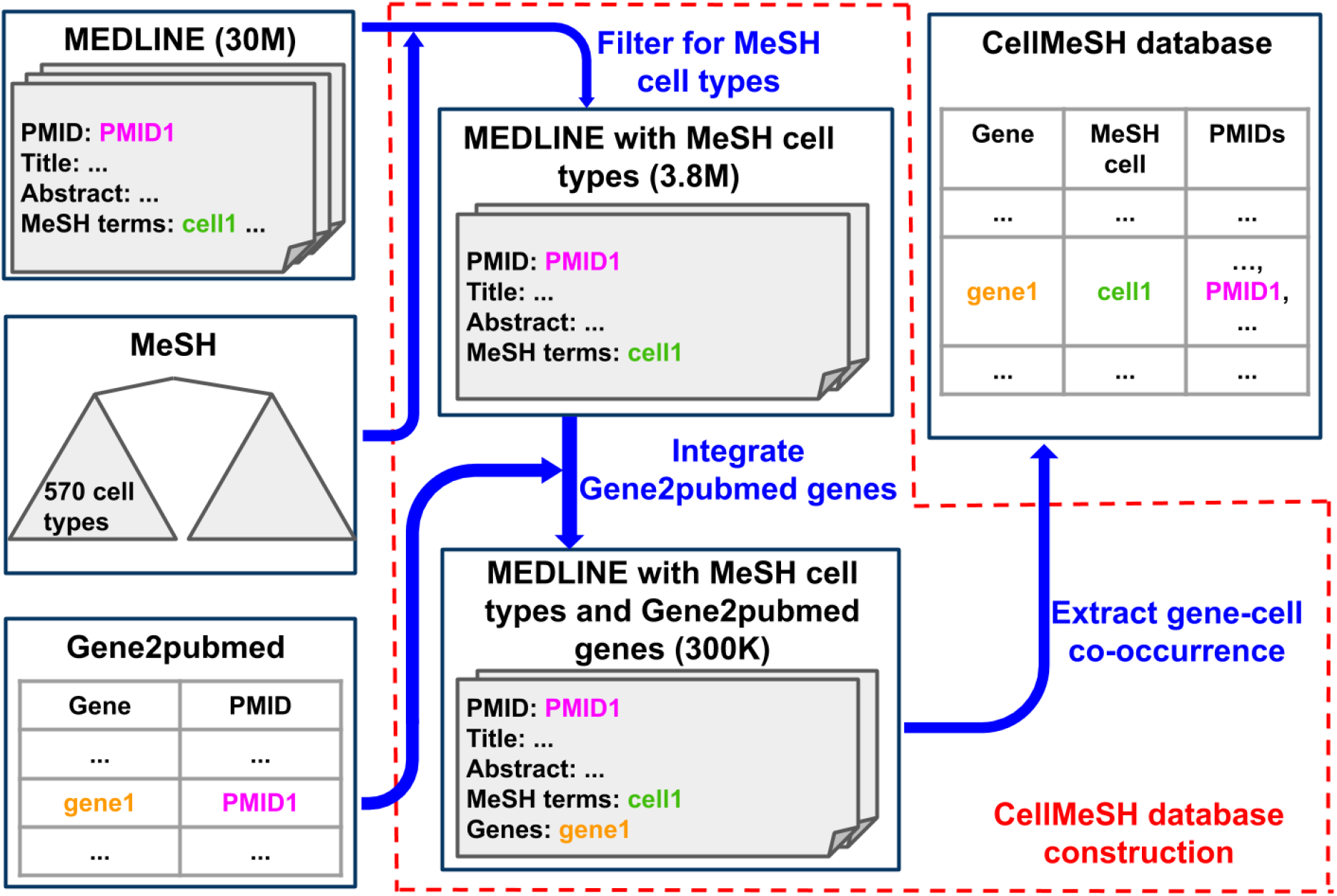
CellMeSH Database Construction. The construction is species-specific. Here we illustrate the construction for the human database. We start with the 30 million MEDLINE references, and keep the ones containing MeSH cell types (there are 3.8 million such references). We further filter away the references not having human genes in Gene2pubmed, after which 300 thousand MEDLINE references remain. Each remaining MEDLINE reference *p* contains several MeSH cell types {c} and several genes {*g*}, and we append *p* to each (*g,c*) pair in the final CellMeSH database.

Next, we integrate gene information from Gene2pubmed [50], a database that links standardized NCBI genes [50] with PubMed articles. Gene2pubmed currently references 20,164 human and 27,322 mouse genes. We construct 2 distinct databases, one for each species - human and mouse. A gene and a cell type are considered to co-occur if there is at least one article that is associated with the cell type in MEDLINE and with the gene in Gene2pubmed. We construct a matrix where each gene is a row and each cell type is a column and the entry denotes the number of articles in which the gene co-occurs with the cell type.

The CellMeSH database statistics are as follows. For human, 3.8% of all possible (20,164×570) genecell pairs have non-zero counts, and around 300,000 PubMed articles each contain at least one pair. For mouse, 2.4% (27,322×570) gene-cell pairs have nonzero counts, and around 209,000 PubMed articles each containing at least one pair.

### Probabilistic query method

There are two major issues with using a literature-derived database. The first issue is publication bias. Some genes or cell types are studied much more than others and, consequently, there are more publications and thus more associations containing those genes or cell types. The second issue is noise in the gene-cell type mapping. The CellMeSH database is inherently noisy, as it links genes and cell types at an article level, and the simple fact of an article mentioning a cell type and a gene together does not imply that the gene serves as a marker for the cell type. This leads to potentially spurious associations between genes and cell types.

First, we highlight how to address the issue of publication bias by applying TF-IDF (Term Frequency-Inverse Document Frequency) [52] which is a reweighting method commonly used in Natural Language Processing [53, 54], and by applying column normalization. Specifically let *w_C_*(*g*) denote the weight, which is the number of co-occurrences of gene *g* in the cell type *C*. Using TF-IDF transformation, the new weight is given by 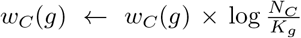 where *N_C_* is the total number of available cell types in the database, and *K_g_* is the total number of cell types with non-zero weights for gene *g*, i.e. 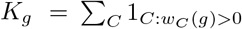. TF-IDF addresses the publication bias of genes since the transformation appropriately deweights commonly occuring genes (since, for these genes, *K_g_* is larger). After TF-IDF transformation, the weight is further adjusted by column normalization: 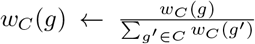. Column normalization addresses the publication bias of cell types since the transformation appropriately deweights the genes occurring in common cell types.

We then query the weight-adjusted database using a probabilistic method, which is designed to address the issue of noise from spurious gene-cell associations. Our query method takes input of *w_C_*(*g*) which is the adjusted weight of gene *g* in cell type *C*. The method also takes input of a query *Q* which is a list of genes. The method outputs the database cell types sorted by their significance to the query.

Our probabilistic query method assumes the following generative model for the observed query data (based on which the inference is performed): (1) a cell type is first chosen (with a uniform prior probability) (2) associated with the cell type is a probability distribution on the genes given by *p*(*g*|*C*). A natural model for *p*(*g*|*C*) is to take it to be proportional to the weight *w_C_*(*g*). (3) However, the previous model ensures that only genes with non-zero weight for the cell type will be present in the query - this need not be the case in our noisy dataset. To model this noise, we assume that with probability 1 – *α* the gene is sampled randomly from the list of all genes, and with probability *α* it is sampled from the cell-type specific gene distribution (in experiments, *α* is fixed as 0.5).

We also denote the total number of genes as *N_g_* and the total number of genes with non-zero weight in cell type *C* as 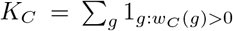. Thus the probability of picking a gene from a cell type can be written as follows:

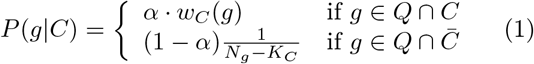

We denote by *P*(*Q*|*C*), the probability that the list of query genes is obtained from a particular cell type *C*. We utilize *P*(*g*|*C*) as the probability that we see gene g in the query given the cell type is *C*. Assuming that each gene is sampled independently, we have 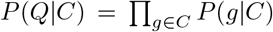. We can then utilize maximum likelihood estimation to predict the cell type *Ĉ*\* that maximizes our chance of seeing the query:

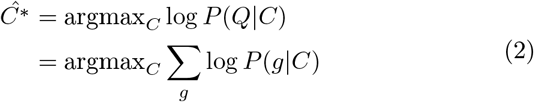

### Cell-type annotation performance

We quantified the cell-type identification performance of CellMeSH for four scRNA-seq datasets with known cell types: the human Peripheral Blood Mononuclear Cells (PBMC) dataset [4], two Tabula Muris (TM) datasets [55], and the Mouse Cell Atlas (MCA) dataset [16].

In our evaluation, we used clusters and reference annotations obtained in the original papers. For each cluster, we extracted the top *n* = 50 differentially expressed genes by 1-vs-rest gene expression ratio. These genes are assumed to be marker genes of the reference cell type and are used as a query input for CellMeSH. We then queried CellMeSH with marker genes for each cluster and visualized results using heatmaps that show how well the top-three retrieved candidate MeSH cell types agree with the reference cell type, according to the mappings between the reference cell types and their correct MeSH cell types, which have been manually made (Additional file 2).

#### Peripheral Blood Mononuclear Cells dataset

The Peripheral Blood Mononuclear Cells (PBMC) dataset is a subset of the data from [4], consisting of 94655 cells and 10 annotated cell types. Cells were originally flow-sorted based on known markers for the corresponding cell type.

Fig. 3 (a) shows the annotation heatmap for the PBMC dataset. The cell types along the y-axis represent the reference query cell types ({*r*}) and those along the x-axis represent a subset of all the candidate MeSH cell types ({*c*}). For a particular query *r*, the heatmap entry (*x* = *c,y* = *r*) is highlighted with a black border, if the candidate *c* matches *r*. A heatmap entry is colored red, if the candidate *c* has rank-1 among the retrieved results for the query *r*, yellow, if it has rank-2 and blue, if it has rank-3. Lighter hues of red, yellow or blue imply that *c* does not match the expected cell type.

**Figure 3.**
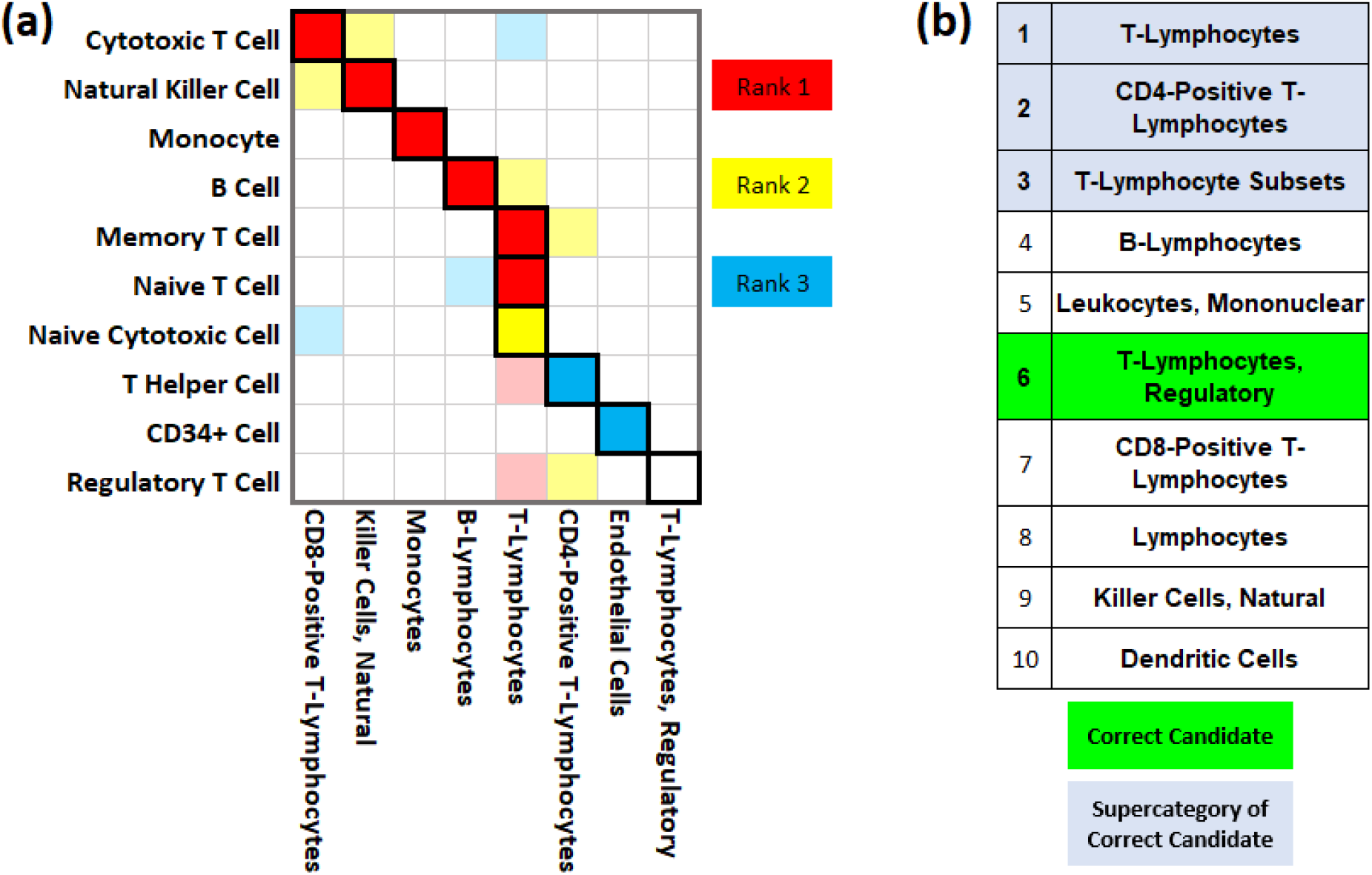
Annotation Results for Human PBMC Dataset. **(a)** is the annotation heatmap where y-axis represents the query cells {*r*} and x-axis represents the candidate cells {*c*}. A border-box for entry (*x* = *c,y* = *r*) indicates *c* is the correct candidate for query *r*. The red, yellow or blue color indicates *c* has rank 1, rank 2, or rank 3 among retrieved results for *r*; the colors are shown lighter if *c* is not the correct candidate. For example, for the query cell B Cell, the red border-box at “B-Lymphocytes” indicates “B-Lymphocytes” is rank 1 in the retrieved results, and it is also the correct result. **(b)** shows the top 10 results for the query Regulatory T Cell, which has an uncolored border-box in the heatmap. This is because its correct candidate cell “T-Lymphocytes, Regulatory” is not among the top 3 (instead it is rank 6) in the retrieved results. In fact the top 3 results are closely related to the correct candidate as they are supercategories of the correct candidate in the MeSH tree hierarchy.This shows that even in cases where our retrieved cell type is not exactly matching the correct cell type, CellMeSH returns reasonable results.

Overall, annotations are accurate as shown by the strong signal along the diagonal. Even where there appears to be no signal on the diagonal, we see that CellMeSH returns reasonable results. For example, the query Regulatory T Cell is not colored at the correct candidate “T-Lymphocytes, Regulatory” because “T-Lymphocytes, Regulatory” is not among the top 3 retrieved results. Fig. 3 (b) indicates that “T-Lymphocytes, Regulatory” is actually rank-6. However, the top 3 results are still promising because they are closely related to “T-Lymphocytes, Regulatory” in the category hierarchy of the MeSH tree. Specifically, “T-Lymphocytes” is a supercategory of “CD4Positive T-Lymphocytes”, which is in turn a supercategory of “T-Lymphocytes, Regulatory”. In addition, “T-Lymphocytes” is also a supercategory of “T-Lymphocyte Subsets”, which is in turn a supercategory of “T-Lymphocytes, Regulatory”.

#### Tabula Muris datasets

For a second example, we turned to the Tabula Muris (TM) dataset [55]. This dataset contains cells that were captured from 20 different tissues in 3-month-old mice and that were clustered into 99 annotated cell types. Compared to the PBMC dataset with only 10 annotated cell types, this dataset is therefore more challenging. The TM dataset contains two subdatasets, with cells captured by using a microfluidic-droplet method (denoted as the TM-Droplet dataset, containing 55656 cells) or by cell sorting (denoted as the TM-FACS dataset, containing 44949 cells). Here we focus on the TM-Droplet dataset (Fig. 4) but the results for TM-FACS are similar (Additional file 3 Fig. 7).

**Figure 4.**
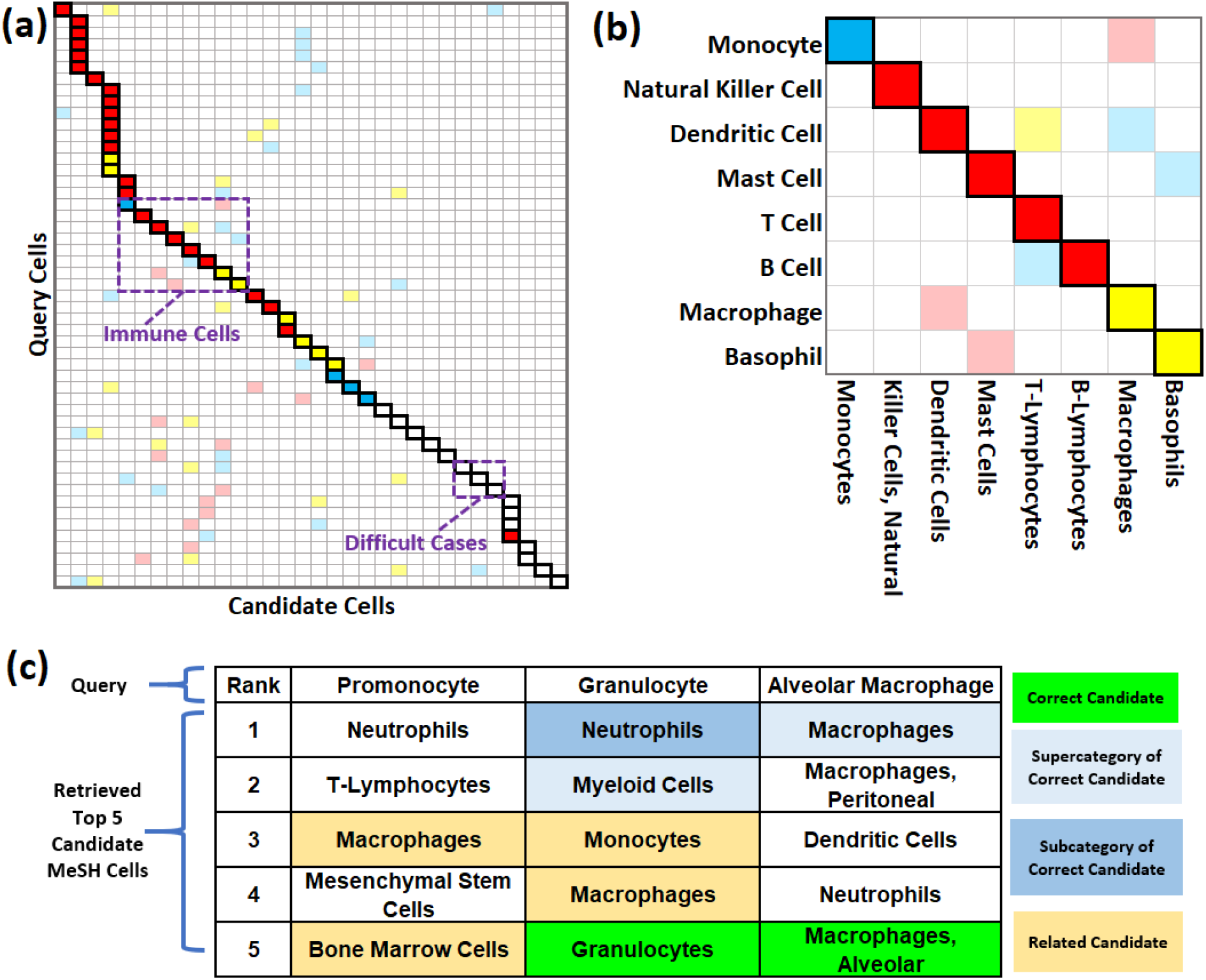
Annotation Results for Tabula Muris Droplet Dataset. **(a)** is the annotation heatmap for queries of all cells. The y-axis represents the query cells {*r*} and x-axis represents the candidate cells {*c*}. A border-box for entry (*x* = *c,y* = *r*) indicates *c* is the correct candidate for query *r*. The red, yellow or blue color indicates *c* has rank 1, rank 2, or rank 3 among retrieved results for *r*; the colors are shown lighter if *c* is not the correct candidate. **(b)** is the annotation heatmap for queries of immune cells, all of which have their correct candidate cells within top 3 retrieved results. **(c)** shows the top 5 results of several query cells corresponding to the uncolored border-boxes in the heatmap. For instance, the query Granulocyte has its correct candidate “Granulocytes” ranked 5-th among the retrieved results. The top 2 results “Neutrophils” and “Myeloid Cells” are accurate to a certain extent because they are the subcategory and supercategory of “Granulocytes” in the MeSH tree. This again shows that in cases where the top retrieved results do not contain the candidate that matches the query exactly, CellMeSH returns reasonable results.

Annotation results for the entire TM-Droplet dataset are summarized in the heatmap shown in Fig. 4 (a) (see Additional file 3 Fig. 6 for detailed cell type names). The diagonal bordered-boxes, indicating the expected annotations, are mostly filled with red, yellow or blue colors used to highlight the top 3 retrievals, which clearly demonstrates the effective annotation ability of CellMeSH. To see this more clearly, in Fig. 4 (b) we focus on the annotation heatmap for only the immune cell types, where all query cells get their correct annotations within top 3 candidates.

There are bordered-boxes forming vertical trajectories in Fig. 4 (a). This is because we manually map several true cell types to the same MeSH cell term due to the limited resolution of the MeSH cell types. For e.g., Luminal Epithelial Cell of Mammary Gland, Kidney Collecting Duct Epithelial Cell, Bladder Urothelial Cell are mapped to “Epithelial Cells”.

There are uncolored bordered boxes in Fig. 4 (a), implying the correct candidate may not exist due to a limit in the coverage of the MeSH cell types, or, more likely, that it is not within the top 3 retrieved results due to the noise in CellMeSH database. Still, the CellMeSH query results provide useful insights into the true cell types, as illustrated in Fig. 4 (c). For instance, the query Promonocyte does not have an exact same MeSH term; the closest term we could manually match is “Monocyte-Macrophage Precursor Cells” (see Additional file 2). However among the top 5 retrieved results, “Monocytes” and “Bone Marrow Cells” are both closely relevant because a Promonocyte is a cell arising from a Monoblast (in Bone Marrow [56]) and developing into a Monocyte [57]. For the query Granulocyte, the correct MeSH term “Granulocytes” is rank-5 in the retrieved results probably due to relatively fewer citations in the database. However the top 2 results “Neutrophils” and “Myeloid Cells” are respectively the subcategory and supercategory of “Granulocytes” in the MeSH tree. “Monocytes” and “Macrophages” are also related as Neutrophils can secrete products that stimulate Monocytes and Macrophages [58]. Similarly, for the query Alveolar Macrophage, the correct MeSH candidate “Macrophages, Alveolar” actually ranks in the top 5. The top rank result “Macrophages” is also close as it is a supercategory of “Macrophages, Alveolar”. See Additional file 4 for more examples regarding to the uncolored bordered-boxes in Fig. 4 (a).

#### Mouse Cell Atlas dataset

The Mouse Cell Atlas (MCA) dataset [16] consists of scRNA-seq data from 6- to 10-week-old mice, sampled from a large variety of tissues. It contains over 200,000 cells after batch effect filtering, and 840 annotated cell types.

Compared to the Tabula Muris dataset, the annotated MCA cell types are much more fine-grained because they each contain additional gene and tissue information (e.g. *r*1 =Alveolar Macrophage_Ear2 high (Lung), *r*2 =Alveolar Macrophage_Pclaf high (Lung)). For our analysis, we collapsed these cell types into 204 cell types based on the cell-type names (e.g. the collapsed *r* =Alveolar Macrophage and its marker genes are obtained from *r*1, *r*2). The MCA heatmap result is generally similar to the PBMC and TM heatmaps, with mostly accurate annotations (Additional file 3 Fig. 8). We also confirmed that using the collapsed 204 cell types shows an overall similar performance as using the original 840 cell types (see Additional file 3 Fig. 5).

**Figure 5.**
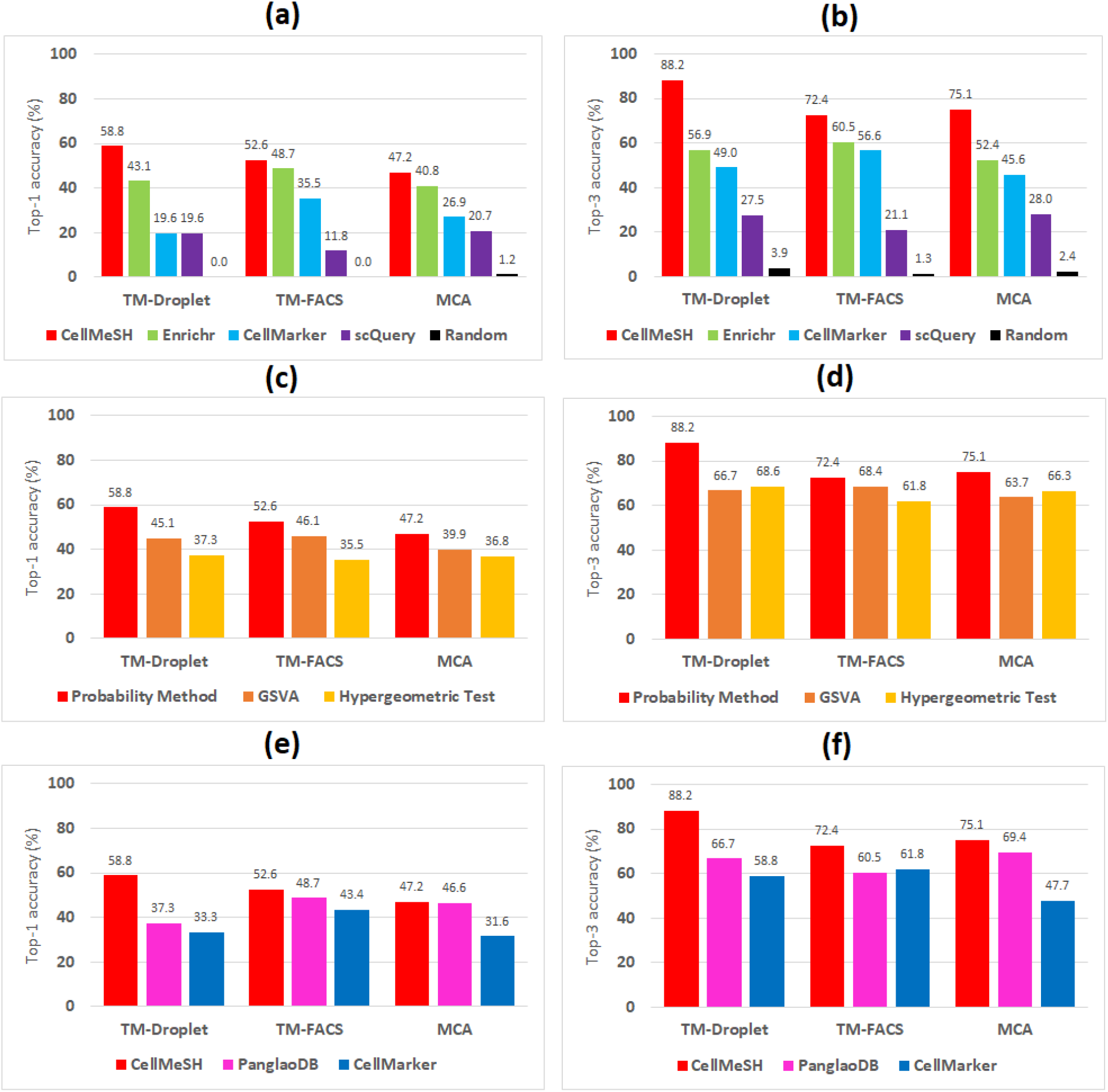
Comparison of CellMeSH to other methods. Each bar plot has y-axis as the top-k (k=1 or 3) accuracy (%) and is grouped by different mouse datasets, and for each group, we show the top-k accuracy of different methods. Top-k accuracy refers to the percentage of queries where one of the candidate cells among the top k retrieved cells is accurate. **(a)(b)** CellMeSH is compared to other systems. For the existing database of CellMarker, we query it using the existing query method of hypergeometric test. **(c)(d)** Probabilistic query method is compared to other query methods. We fix CellMeSH database to be queried by all query methods. **(e)(f)** CellMeSH database is compared to other databases. Note that CellMeSH and CellMarker are both non-binary gene-cell matrices and therefore we use probabilistic method to query,whereas PanglaoDB is a binary gene-cell matrix and we use hypergeometric test to query. CellMeSH demonstrates a consistent better top-1 and top-3 accuracy than other methods for all datasets.

### Performance of top-k accuracy

We compared CellMeSH to several existing methods [20, 22–25, 43] (Table 1) by using top-k (*k* = 1, 3) accuracy for the three mouse datasets (TM-Droplet, TM-FACS and MCA). We use the mouse datasets instead of the human PBMC dataset, because they contain many more queries (51, 76 and 191 queries^[1]^ respectively) than the PBMC dataset (only 10 queries), and therefore can show more reliable quantitative trends.

We queried CellMeSH with previously extracted marker genes for each cluster, and calculated the top-k accuracy (percentage of queries that retrieve the expected candidate cell type in the top k results) for each dataset. We similarly queried existing methods and obtained top-k results for each dataset. The query results of existing methods could derive from a different ontology and therefore contain different cell type names from the MeSH terms. In order to calculate the accuracy of these methods, we thus manually created mappings between the given query cell types and the candidate cell types from other ontologies, as summarized in Additional file 2.

#### Overall top-k accuracy gain

In Fig. 5 (a)(b), we first compare CellMeSH with two other web servers: Enrichr [20] and scQuery [43] as they share the most similar input and output formats (Table 1). We are unable to compare to the web servers of PanglaoDB [25], CellMarker [22] and CellFinder [21] in an automatic way. Instead, we downloaded the existing CellMarker [22] database and queried it using the hypergeometric test. We also include the random retrieval results as a lower bound.

All of the methods are significantly better than random guessing, and CellMeSH provides the most accurate results for all three mouse datasets. Specifically, in the TM-Droplet dataset, CellMeSH achieved top-1 accuracy of 58.8%, meaning that in 58.8% of queries, the first retrieved candidate cell type is correct. The top-1 accuracy is 15.7% higher than that of the second best method, Enrichr. This is to be expected because the Enrichr cell types essentially come from the Mouse Gene Atlas (MGA) database [59], which contains only 96 cell types. Besides, some of the MGA cell types (such as “Heart”, “Kidney”, and “Stomach” etc) actually refer to organs. We find that CellMeSH has higher coverage and resolution than the other methods including Enrichr. For example, for query Classical Monocyte, while CellMeSH returns “Monocytes” as the first candidate, there is no monocyte term covered in MGA and Enrichr returns its first result as “Macrophage Bone Marrow 6hr LPS”. For query Duct Epithelial Cell, while CellMeSH returns “Epithelial Cells” as the first result, Enrichr returns the organ terms “Bladder”, “Liver” and “Stomach” as its top 3 results (see Additional file 5 for details). The top-3 accuracy of CellMeSH further increases to 88.2% (this implies that 88.2% of queries get at least one of the top 3 results correct), which is 31.3% higher than that of Enrichr.

CellMeSH also consistently outperforms other methods on the other two datasets. Its top-1 (or top-3) accuracy is 3.9% (or 11.9%) higher in the TM-FACS dataset, and 6.4% (or 22.7%) higher in the MCA dataset, than the second best method, Enrichr.

#### Top-k accuracy gain from probabilistic method

Both the CellMeSH database and the probabilistic query method contribute to the overall top-k accuracy gains of CellMeSH. To isolate the contribution of the probabilistic query method to the overall CellMeSH performance, we compared it to the more established hypergeometric test [23] and GSVA [24] that are suggested in [60], by querying the same CellMeSH database for the three mouse datasets.

As shown in Fig. 5 (c)(d), the probabilistic method performs uniformly better than other methods. Compared to the best performance of GSVA and the hypergeometric test, the probabilistic query method has a top-1 accuracy gain of 13.7%, 6.6% and 7.3% in the TM-Droplet, TM-FACS and MCA datasets respectively. The numbers for top-3 accuracy gain are 19.6%, 3.9% and 8.8%.

#### Top-k accuracy gain from CellMeSH database

To isolate the contribution of the CellMeSH database, here, we compare the performance of alternative databases obtained as follows. We prepared gene-cell co-occurrence matrices by aggregating the cell-type marker-genes files from PanglaoDB [25] and CellMarker [22], both of which are manually compiled from the literature. We then compared the CellMeSH database to these two databases.

We query the CellMeSH and CellMarker databases using the probabilistic query method, since these databases are count-valued matrices, which can be handled effectively by the probabilistic query method, as illustrated in Fig. 5 (c)(d) and in Additional file 3 Fig. 4. For PanglaoDB, the query method is the hypergeometric test, since the database is essentially a binary matrix.

As Fig. 5 (e)(f) illustrates, using the CellMeSH database achieves a higher accuracy than using the PanglaoDB and CellMarker databases. Compared to the best performance out of PanglaoDB and CellMarker databases, the CellMeSH database has top-1 accuracy gain of 21.6%, 3.9% and 0.6% in the TM-Droplet, TM-FACS and MCA datasets respectively. The numbers for top-3 accuracy gain increase to 21.6%, 10.5% and 5.7%.

#### Impact of gene number

Our evaluations so far have been using the fixed top *n* =50 differentially expressed genes as the marker genes for each query cell type. In addition, here the performance tends to peak around n = 50 for most of the methods (see Additional file 3 Fig. 1 to 3); A smaller number of genes may not provide sufficient information, and a larger number of genes may bring more noise, both of which could result in degraded annotation performance. If we select the optimal number of genes for each method (e.g. different settings of the database and the query method), the CellMeSH database together with the probabilistic query method still consistently outperforms all other methods for all of the datasets (see Additional file 5).

### CellMeSH web server and API

CellMeSH has a stand-alone web server!^[2]^, which is able to take in a list of marker genes and returns a ranked list of predicted MeSH cell types, together with the supporting genes and PubMed articles for further reference (Fig. 1). The web server also provides options to use other databases (e.g. CellMarker) and query methods (e.g. GSVA, hypergeometric test).

We have open-sourced the CellMeSH database and the probabilistic query method^[3]^ as Python API to assist the community efforts on automating the cell type identification. We have integrated it into our recently developed web server UNCURL-App [28] for interactive scRNA-seq data analysis, which combines data preprocessing, dimensionality reduction, clustering, differential expression, cell-type annotation and interactive re-clustering into an online graphical user interface.

## Discussion

We have developed CellMeSH, a method with accompanying web server and API to identify cell types directly from literature, in order to make the scRNA-seq analysis more convenient. Experiments on both human and mouse scRNA-seq datasets demonstrate CellMeSH’s superior cell-type identification performance.

Nevertheless, there are still several limitations with CellMeSH. Particularly, the cell-type annotations provided by MeSH terms are somewhat coarse, and might not be enough to represent a comprehensive listing of all fine-grained (sub) cell types present in model organisms such as human or mouse. Moreover, for other species, even other model organisms, gene-cell information could be limited due to a lack of indexed publications.

One way the CellMeSH database can be extended is through natural language processing on full text articles (including supplementary files). Ideally such an approach would enable the identification of new cell types in papers using unsupervised or semi-supervised named entity recognition [61, 62].

There are also terms within the MeSH ontology that may be useful but are not under the “Cell” heading, such as tissues, organs, and diseases. Designing the query methods utilizing these information is an interesting future direction. For instance, we can refine our search scope if we know the tissue information of the query; or if such information is missing we could provide them from an extended Cell/Tissue-MeSH database.

## Methods

### Experiment I/O

For each dataset (PBMC, TM Droplet, TM-FACS and MCA), we have prepared the set of queries denoted as *D* = {(*r_i_,Q_i_*)}, *i* = 1,…,*N_D_*, where the *i*-th query has reference cell type *r_i_* and the query genes are *Q_i_* (e.g. *Q_i_* contains top *n* = 50 differentially expressed genes). There are *N_D_* queries for dataset *D*.

We queried different methods including external web server systems and combinations of different query methods and databases (Fig. 5). Generally, for a particular query (*r_i_, Q_i_*), we can represent the query results as a list of ranked candidate cell types *C_i_* = {*c*_*i*,1_, *c*_*i*,2_,…, *c_i,M_*} where *M* is the total number of candidate cell types. Depending on the systems (Enrichr, scQuery) or the databases (CellMarker, PanglaoDB) used, Ci differs in terms of candidate cell names and length *M*. As the candidate cell type (e.g. *c_i,j_* = “Macrophages, Alveolar”) and true label (e.g. *r_i_* = Alveolar Macrophage) may not come from the same ontology and thus have different expressions, their mappings are manually verified, as provided in Additional file 2.

### Heatmap

To obtain the heatmap, we first prepare the heatmap matrix from query results where y-axis represents the reference (true) cell types {*r*} and x-axis as the union of the correct candidate cell types {*c*} (each *c* is a correct candidate for at least one *r*). The matrix has its value at a particular entry (*x* = *c, y* = *r*) equal to *j* where *j* (1-based) refers to the rank of *c* in the retrieved results for *r*. In the heatmap, we highlight the (*c, r*) entries using bordered-boxes if *c* is the correct candidate for *r*, according to our manual mappings (Additional file 2). We also fill the (*c, r*) entries by red, yellow or blue color if *c* has rank-1, rank-2 or rank-3 among the retrieved results for query *r*. To understand these: for a particular query *r*, a red bordered-box indicates *c*, the top 1 retrieval, is the correct result (e.g. “correct hit”); An uncolored bordered-box indicates the correct result of *r* is not among the top 3 retrieved results (e.g. “miss hit”). We tune red, yellow or blue color lighter to further imply *c* is not an expected result of *r* (e.g. “false positive hit”). We reorder the rows and columns of the heatmap so that the bordered-boxes could form a diagonal trajectory of expected annotation results.

### Top-k accuracy

The top-k accuracy (e.g. *k* = 1,3) of the dataset *D* is defined as:

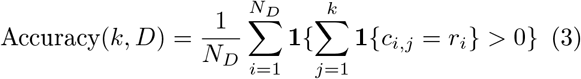

This indicates the ratio of the queries where at least one of the top k retrieved candidate cell types matches the true labels. The justification to use the top-k accuracy is that in practice users will be interested in only a small number of retrieved cell types.

The judgement of whether a candidate cell matches the reference cell is slightly less strict in top-k accuracy than in annotation heatmap, where each query corresponds to only one correct candidate cell type. In top-k accuracy, each query cell may have more than one correct candidate cell types (see Additional file 2). This is because some top retrieved results, though not strictly matching the true label, can still be considered as correct. For example, for the query Granulocyte in Fig. 4 (c), we consider both “Granulocytes” and “Myeloid Cells” as correct retrieved results, in top-k accuracy. We apply this to all of the other methods (e.g. Enrichr, scQuery, CellMarker, and PanglaoDB), actually in a more generous way. For example, for the query Cardiac Muscle Cell, Enrichr returns “Heart” as the top candidate cell, and we also consider it to be correct.

### Hypergeometric test

Hypergeometric test is a simple and effective statistical test that has been widely used in various gene set enrichment analysis [63, 64], to query a set of genes to a noise-free binary database.

In order to use a hypergeometric test to query a weighted database with a set of genes, the database first needs to be binarized [60]. We binarize the noisy CellMeSH database by setting the raw genecell count values in the database to 0 if the count is below a threshold (e.g. 4) and to 1 otherwise. We selected threshold 4 because the corresponding binary CellMeSH database, compared to the binary databases obtained by using other thresholds, achieved the best top-1 accuracy when we queried the database with 335 queries that were prepared from the CellMarker records (Additional file 3 Fig. 9). For the noise-free CellMarker database, the database is binarized using threshold 0. For the PanglaoDB database, it is already binary.

For each query *q* and each candidate *c*, we calculate an enrichment score. Let *n* be the number of genes in *q*. Let *K* be the number of genes co-occurring with *c* in the binary database. Let *N* be the total number of genes. Let *k* be the number of overlapping genes between *q* and *c*. Then, the probability of *k* genes occuring in both query *q* and candidate *c* can be expressed as: 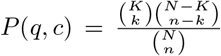. A smaller probability value indicates higher significance for the candidate *c* to be related to the query *q*.

### GSVA

Gene Set Variation Analysis (GSVA) [24] was developed for microarray and bulk RNA-seq analysis, in order to query a database of gene-sets (essentially a noise-free binary database, with rows as genes and columns as pre-defined gene-sets that are related to certain biological activity) with an ordered-list of genes. Instead, in our application, the database can be thought of as an ordered list of genes (for each celltype) and the query is simply a gene-set. We therefore can adapt GSVA for our problem, and we explain the details in the following paragraphs.

Specifically, the original GSVA takes as input, the gene-sample expression matrix and normalizes by using a Gaussian or Poisson kernel estimation [24], to evaluate the relative ordering of genes inside each query sample in the context of the sample population distribution. Each gene then gets its rank in each query sample.

GSVA calculates an enrichment score similar to the Kolmogorov-Smirnov (KS) statistic [65] between each query sample and each candidate gene-set from database, by walking along the ranked genes in the query sample. Specifically, GSVA keeps a running statistic, which it increases if the gene is in the candidate gene-set, and decreases if the gene is not in the candidate gene-set, and retains the maximum deviation of the running statistic as the enrichment score.

In our implementation, since the roles of the query and database are flipped, we normalize the weighted *database* by TF-IDF, to evaluate the relative ordering of genes inside each candidate cell type. Each gene then gets its rank in each candidate cell type (unlike in the original GSVA, where each gene gets a rank for each query sample).

We calculate the enrichment score between each candidate cell from database and each query set from input, by walking along the ranked genes in the candidate cell type. Similarly, the adapted GSVA keeps a running statistic, which it increases if the gene is in the query, and decreases if the gene is not in the query, and retains the maximum deviation of the running statistic as the enrichment score.

In our setting, we found that using TF-IDF normalization instead of Gaussion or Poisson kernel normalization (as suggested in GSVA) improved performance (Additional file 3 Fig. 10 (a)). Furthermore, the adapted GSVA algorithm where the roles of query and database are flipped also provides an improvement for our setting (Additional file 3 Fig. 10 (b)).

### CellMarker database

We constructed the CellMarker database (containing two gene-cell co-occurrence matrice for mouse and human respectively) by aggregating the cell-type markergenes files downloaded from the CellMarker website^[4]^. These files contain species of mouse and human. For mouse, there are 1255 records covering 673 PubMed articles, and most of the records each correspond to a cell type in Cell Ontology [66] term with its marker genes. We have skipped the records that have no marker genes or have non-traditional cell-type names (e.g. “Lee et al.Cell.A:1”). The resulting mouse genecell co-occurrence matrix has 7208 genes and 313 cell terms, where the matrix value for a particular genecell pair represents the number of records (i.e. publications) they co-occur. Similarly for human, there are 2869 records covering 1763 PubMed articles. The resulting human gene-cell co-occurrence matrix has 8973 genes and 364 cell terms. The CellMarker database is mostly sparse, with < 1% positive counts, and maximum counts below 10.

### PanglaoDB database

We constructed the PanglaoDB database (containing one gene-cell binary matrix for both mouse and human) by associating the cell-type marker-genes downloaded from the PanglaoDB website^[5]^. The resulting PanglaoDB database, a binary gene-cell matrix, has 4679 genes (for both mouse and human) and 178 cell types. There are 8230 (1%) non-zero entries.

## Supporting information

Additional file 1 - Query Genes

Additional file 2 - Label Mapping

Additional file 4 - Heatmap Difficult Cell Analysis

Additional file 5 - Top k Retrieval Results

## Ethics approval and consent to participate

Not applicable.

## Consent for publication

Not applicable.

## Availability of data and materials

The datasets supporting the conclusions of this article are all included in additional files.

The CellMeSH webserver is at https://uncurl.cs.washington.edu/db_query.

The CellMeSH Database and Probabilistic Query Method are available at Github site https://github.com/shunfumao/cellmesh under MIT License.

## Author’s contributions

Conceptualized the tool: SM, YZ, GS, SK. Implemented the probabilistic method and database: SM, YZ Implemented the web server: YZ Experiment and analysis: SM, YZ Wrote and edited the paper: SM, YZ, GS, SK.

## Acknowledgements

We’d like to thank Sumit Mukherjee for giving us many insightful suggestions and pointing us to many useful resources.

## Additional Files

Additional file 1 - Query Genes

This file records the top 1000 genes of each query cell for different datasets.

Additional file 2 - Label Mapping

This file lists the mappings between the query cells and their correct candidate cells from CellMeSH, CellMarker, PanglaoDB, Enrichr and scQuery resources, for performance evaluation purpose.

Additional file 3 - Supplementary Figures

This file contains additional figures to support the main text.

Additional file 4 - Heatmap Difficult Cell Analysis

This file contains additional examples to support Fig 3 (b) and Fig 4 (c).

Additional file 5 - Top k Retrieval Results

This file records the top-k accuracy versus the number of genes for different approaches on different datasets. It also includes the detailed top 3 retrieved cell types and their evaluation for various approaches.

## Footnotes

[1] Some queries (e.g. Cell in Cell Cycle) are excluded as there are no matching candidates in any databases (e.g. CellMeSH, CellMarker, PanglaoDB etc).

[2] https://uncurl.cs.washington.edu/db_query

[3] https://github.com/shunfumao/cellmesh

[4] http://biocc.hrbmu.edu.cn/CellMarker/download.jsp

[5] https://panglaodb.se/markers.html

