## Additional file 3 - Supplementary Figures for "CellMeSH: Probabilistic Cell-Type Identification Using Indexed Literature"

May 30, 2020

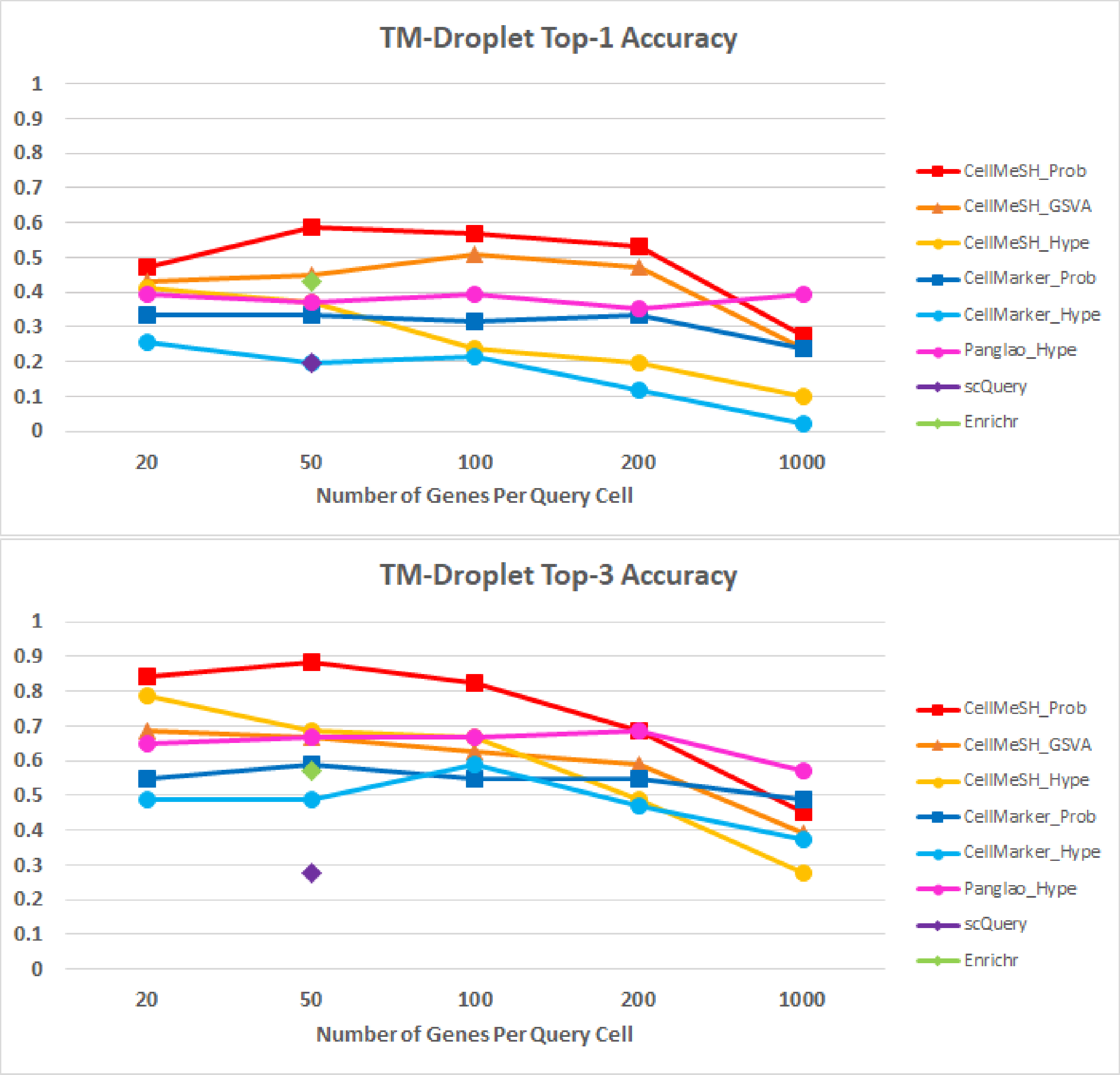


Figure 1: **Top-k Accuracy versus Number of Genes on TM-Droplet Dataset.** Each subplot has its y-axis as the top-k ($k = 1, 3$) accuracy and x-axis as the number of marker genes (e.g. $n$) of each query cell. The curves of different colors correspond to different approaches to identify cell-types. Most approaches peak around $n = 50$.

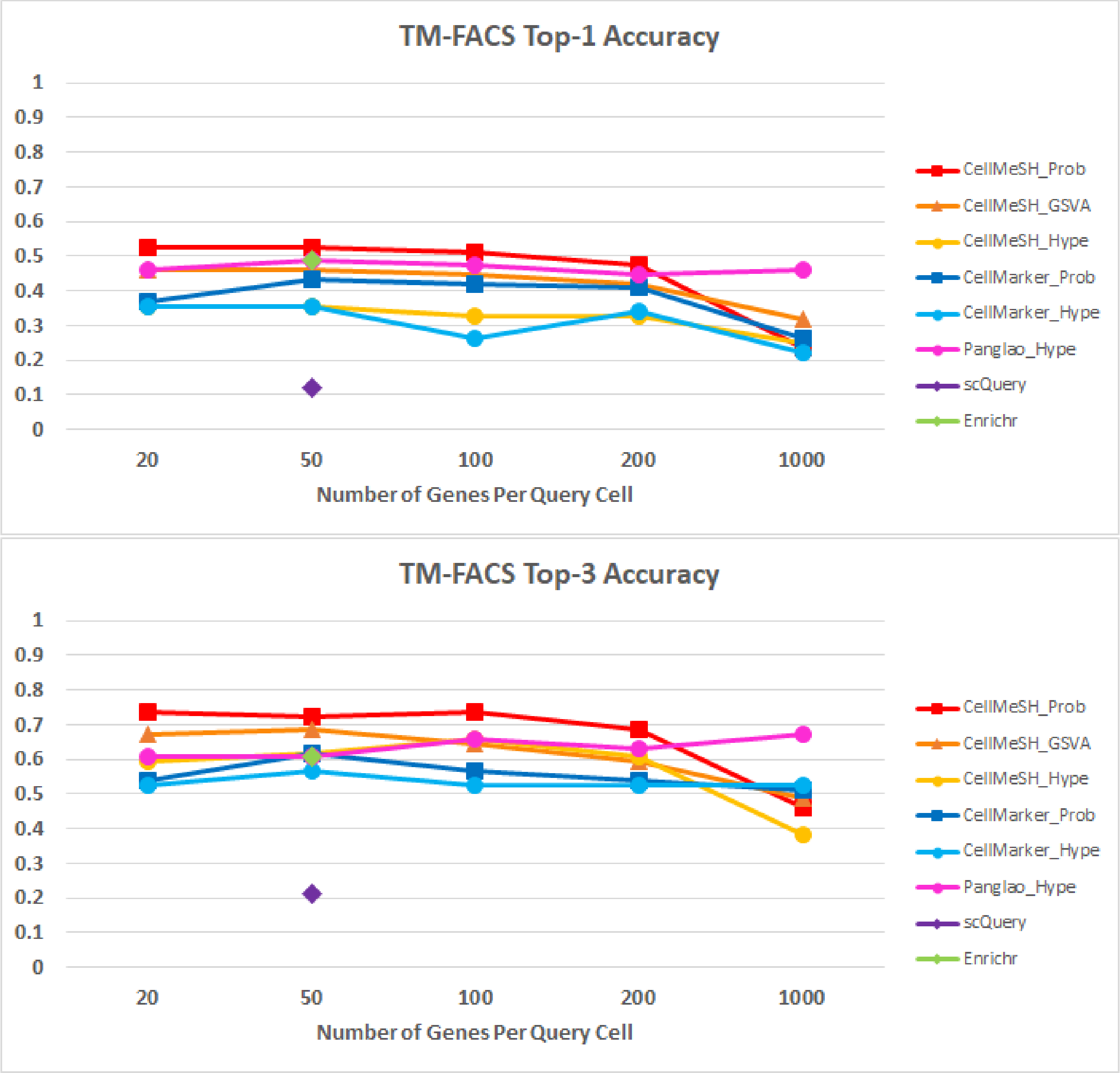


Figure 2: **Top-k Accuracy versus Number of Genes on TM-FACS Dataset.** Each subplot has its y-axis as the top-k ($k = 1, 3$) accuracy and x-axis as the number of marker genes (e.g. $n$) of each query cell. The curves of different colors correspond to different approaches to identify cell-types. Most approaches peak around $n = 50$.

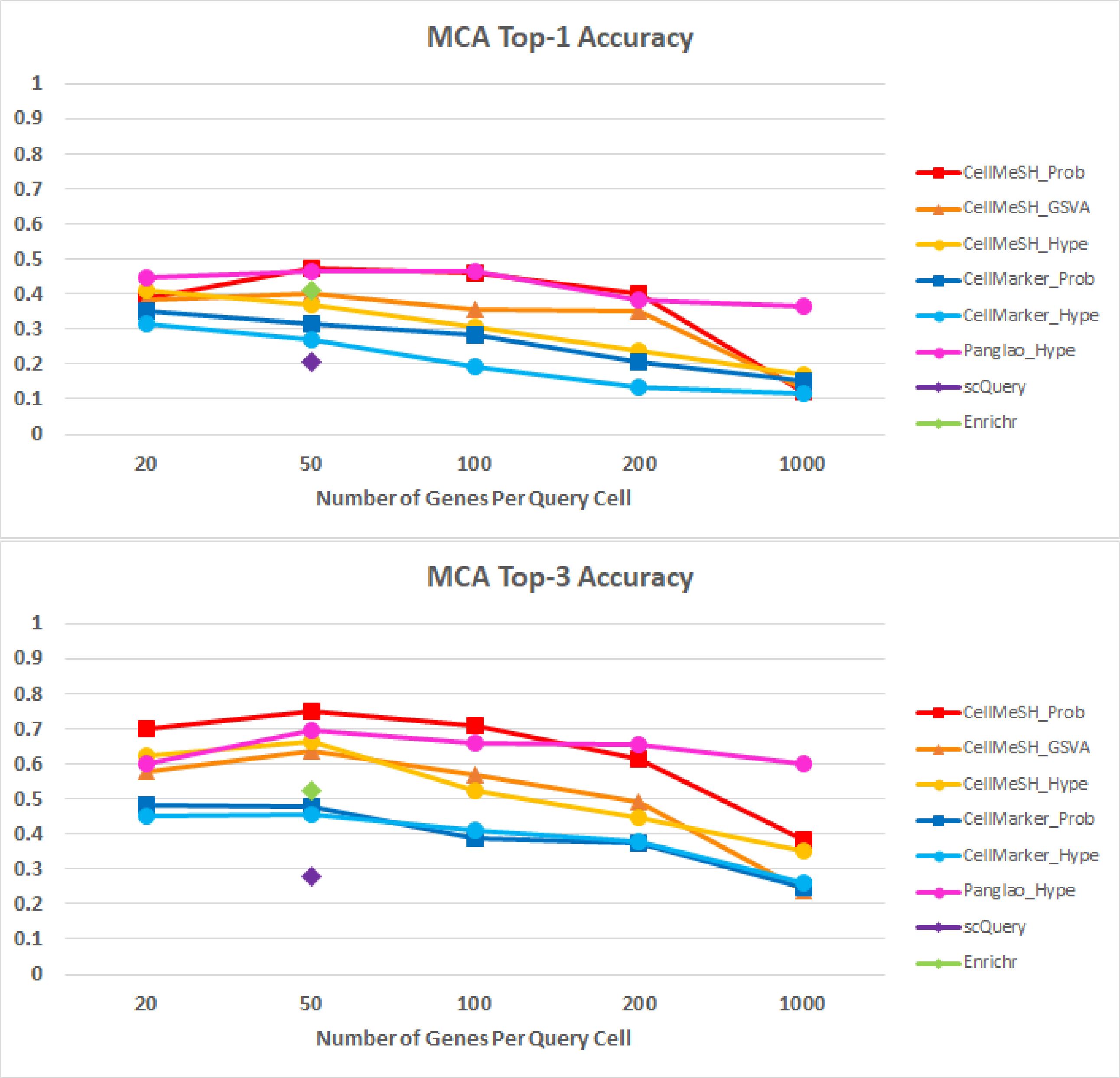


Figure 3: **Top-k Accuracy versus Number of Genes on MCA Dataset.** Each subplot has its y-axis as the top-k ($k = 1, 3$) accuracy and x-axis as the number of marker genes (e.g. $n$) of each query cell. The curves of different colors correspond to different approaches to identify cell-types. Most approaches peak around $n = 50$.

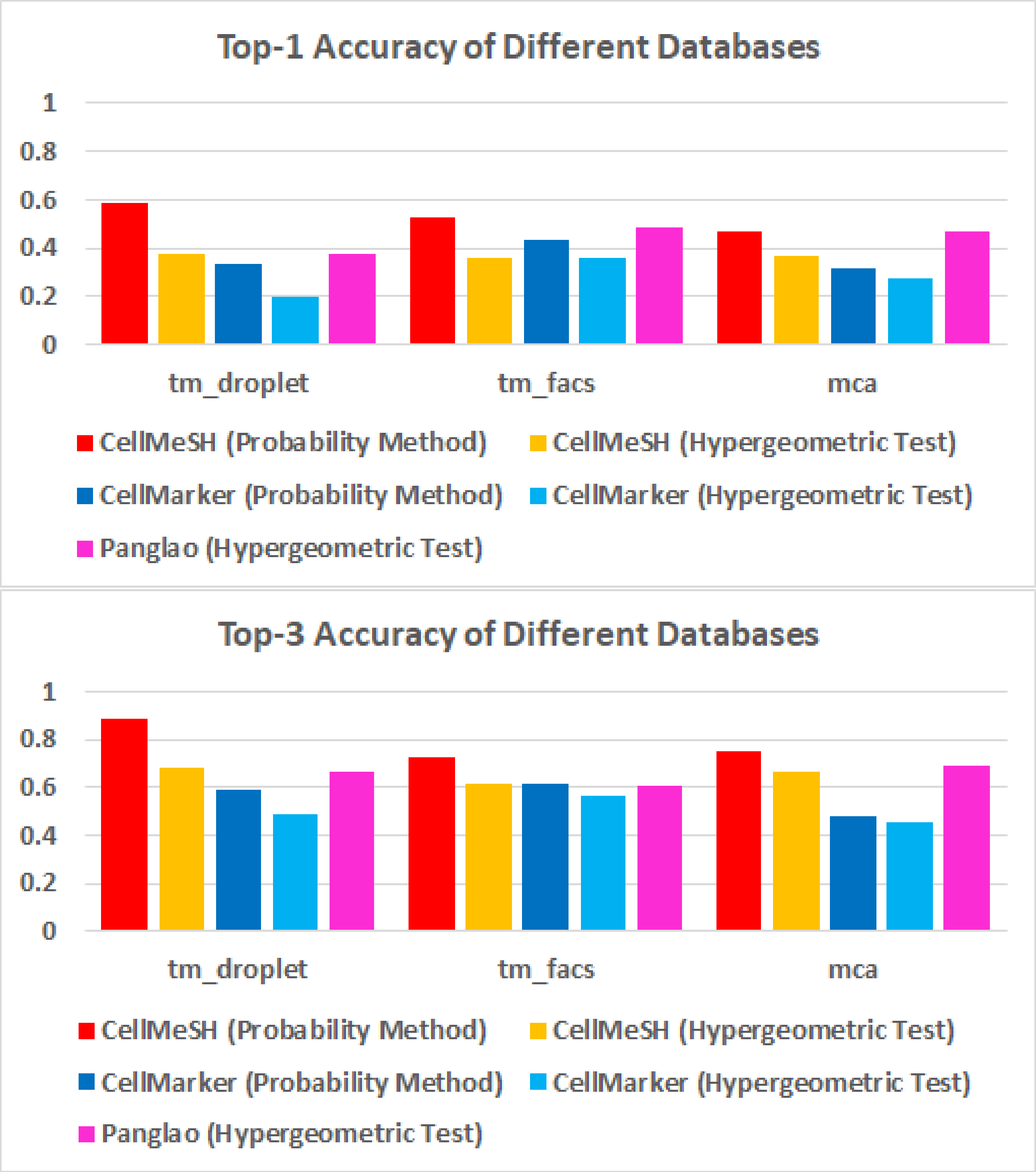


Figure 4: **Top-k Accuracy on Different Databases.** Each subplot has its y-axis as top-k ($k = 1, 3$) accuracy and x-axis as different datasets. Bars of different colors represent different approaches to identify cell-types. When we fix query method as hypergeometric test, PanglaoDB results (pink) actually achieves better performance than CellMeSH (orange) and CellMarker (light blue). As CellMeSH and CellMarker databases are essentially non-binary matrice and contains more noise, using probabilistic query method will better extract the information. CellMeSH (red) eventually performs the best by utilizing the probabilistic query method.

(a) Top-1 Accuracy

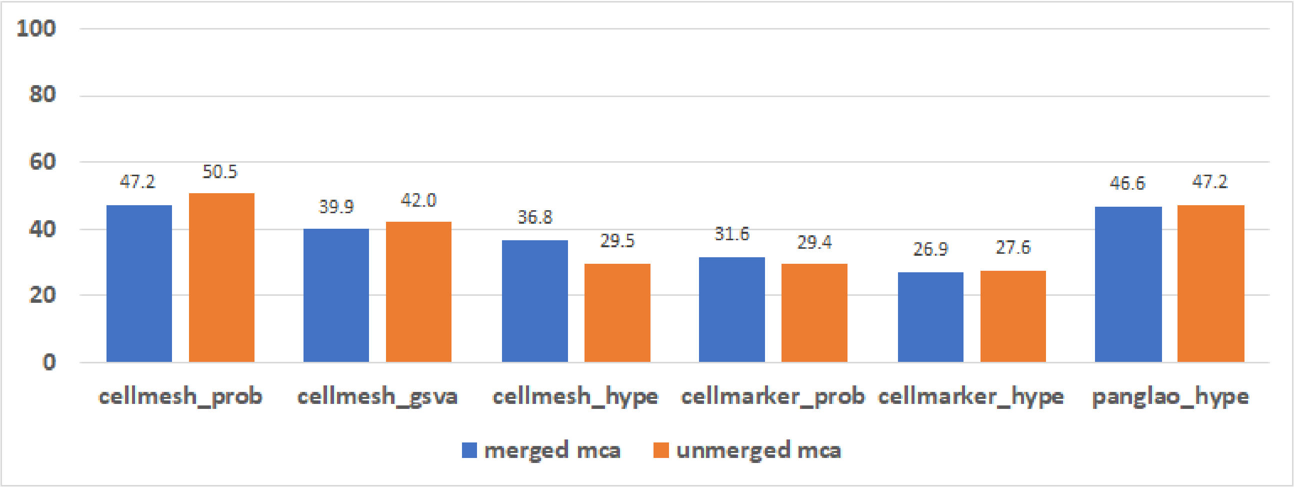


(b) Top-3 Accuracy

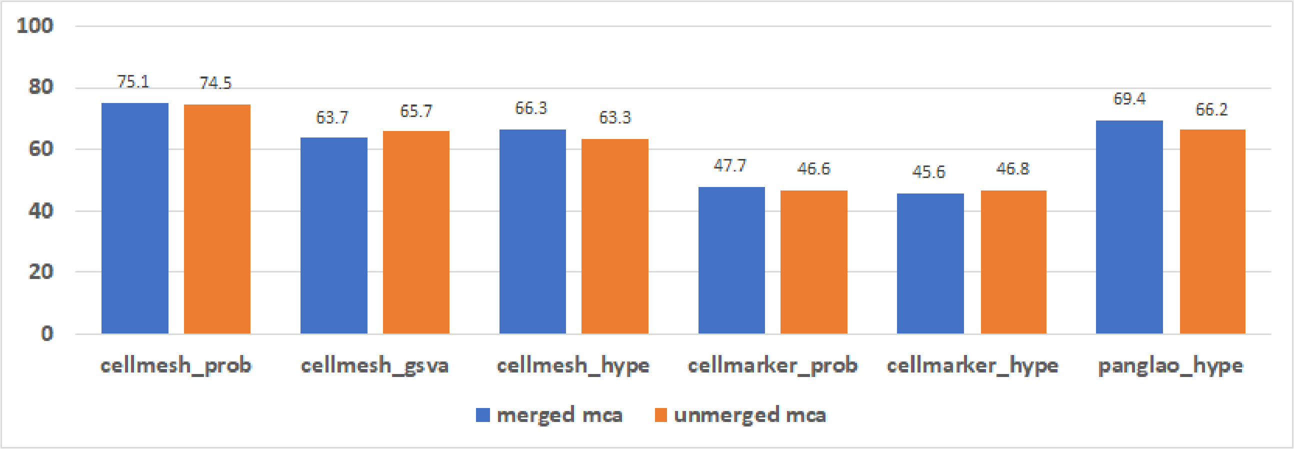


Figure 5: **Compare top-k accuracy using merged and unmerged MCA queries.** We collapsed the original 840 MCA cell-types (unmerged queries; e.g. $r1$ =Alveolar Macrophage_Pclaf high (Lung), $r2$ =Alveolar Macrophage_Ear2 high (Lung)) into the 204 cell-types (merged queries; e.g. $r$ =Alveolar Macrophage). We removed some cell-types which have no equivalent candidate cell-types across any database, resulting 671 unmerged queries and 191 merged queries. (a) compares the merged queries (blue) and the unmerged queries (orange) in terms of top-1 accuracy. The y-axis represents the top-1 accuracy in percentage, and the x-axis represents different combinations of databases (CellMeSH, CellMarker, PanglaoDB) and query methods (Probablistic, GSVA, Hypergeometric test). Using merged queries, CellMeSH (the blue cellmesh_prob bar) achieves 47.2% top-1 accuracy, indicating 47.2% of the queries have their first retrieved result as correct. Overall, the merged queries and unmerged queries have very similar performance. Note that the gain of the best CellMeSH (cellmesh_prob) over the second best PanglaoDB (panglao_hype) is 0.6% (= 47.2% − 46.6%) by using merged queries, which is actually less than 3.3% (= 50.5% − 47.2%) by using unmerged queries. (b) is similar to (a) and compares the merged queries (blue) and the unmerged queries (orange) in terms of top-3 accuracy.

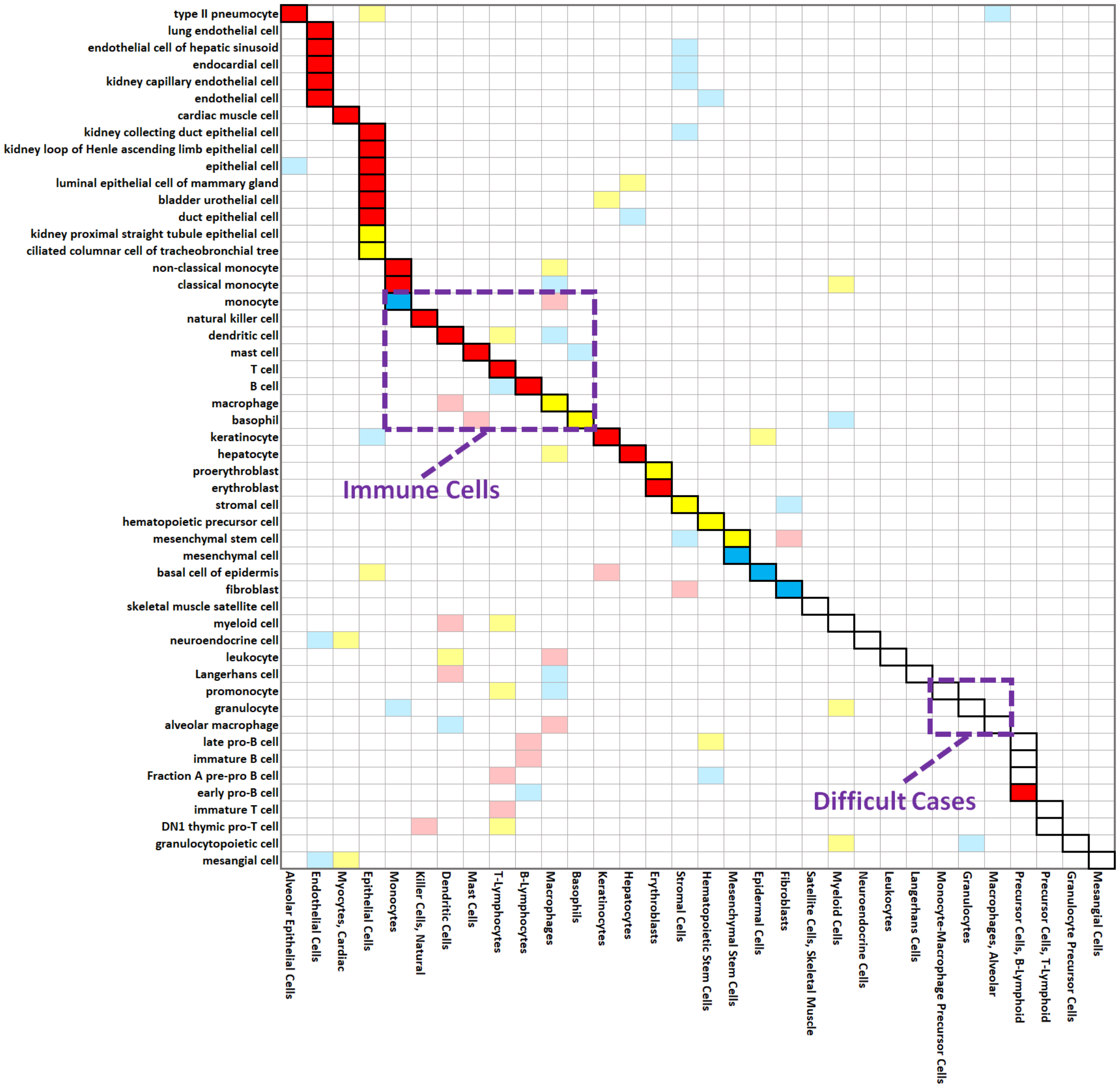


Figure 6: **Annotation Heatmap on TM-Droplet Dataset.** The annotation heatmap has its y-axis representing the query cells $\{r\}$ and x-axis representing the candidate cells $\{c\}$. A border-box for entry $(x = c, y = r)$ indicates $c$ is the correct candidate for query $r$. The red, yellow or blue color indicates $c$ has rank 1, rank 2, or rank 3 among retrieved results for $r$; the colors turn lighter if $c$ is not the correct candidate.

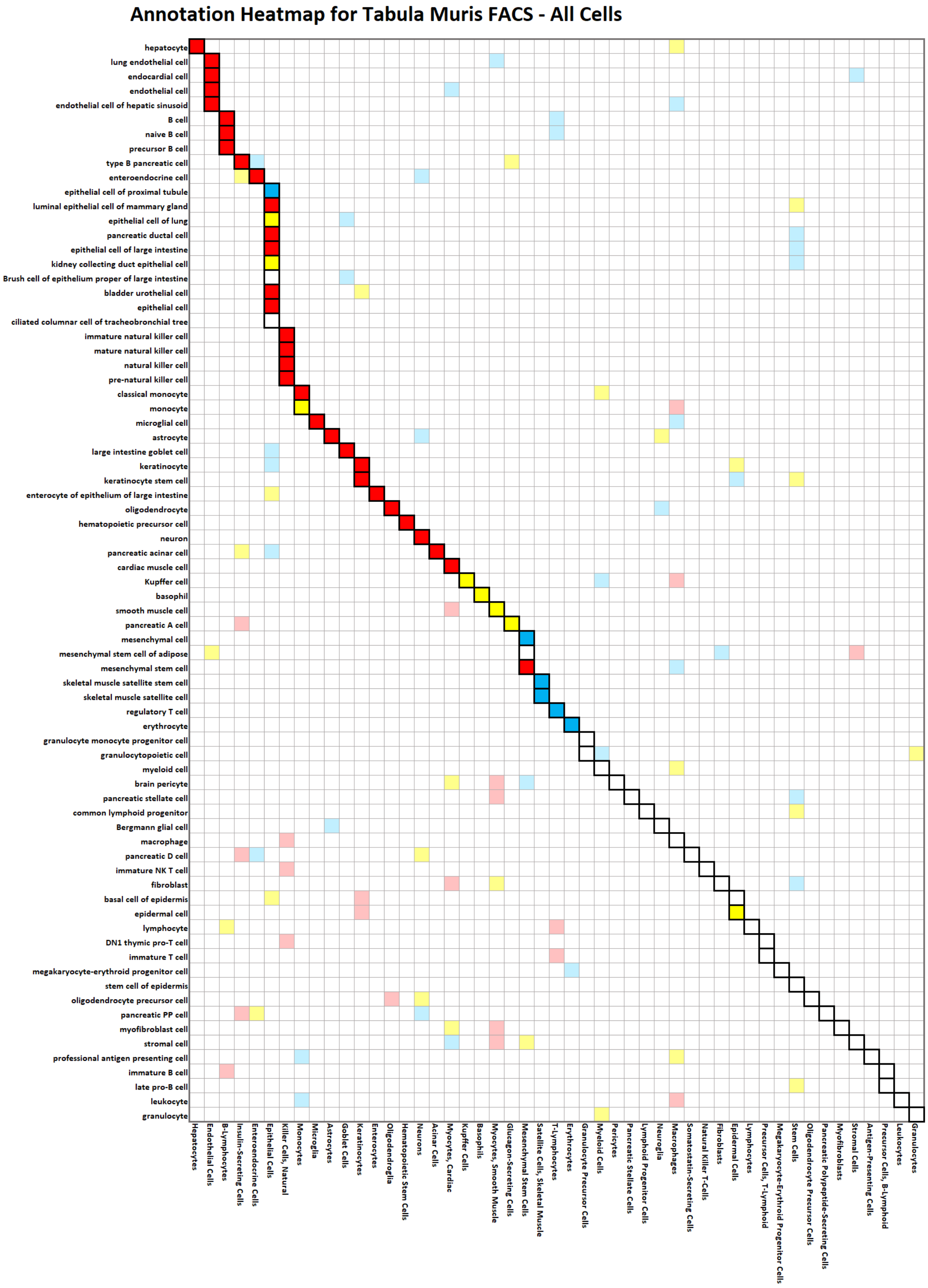


Figure 7: **Annotation Heatmap on TM-FACS Dataset.** The annotation heatmap has its format similar to Fig 6.

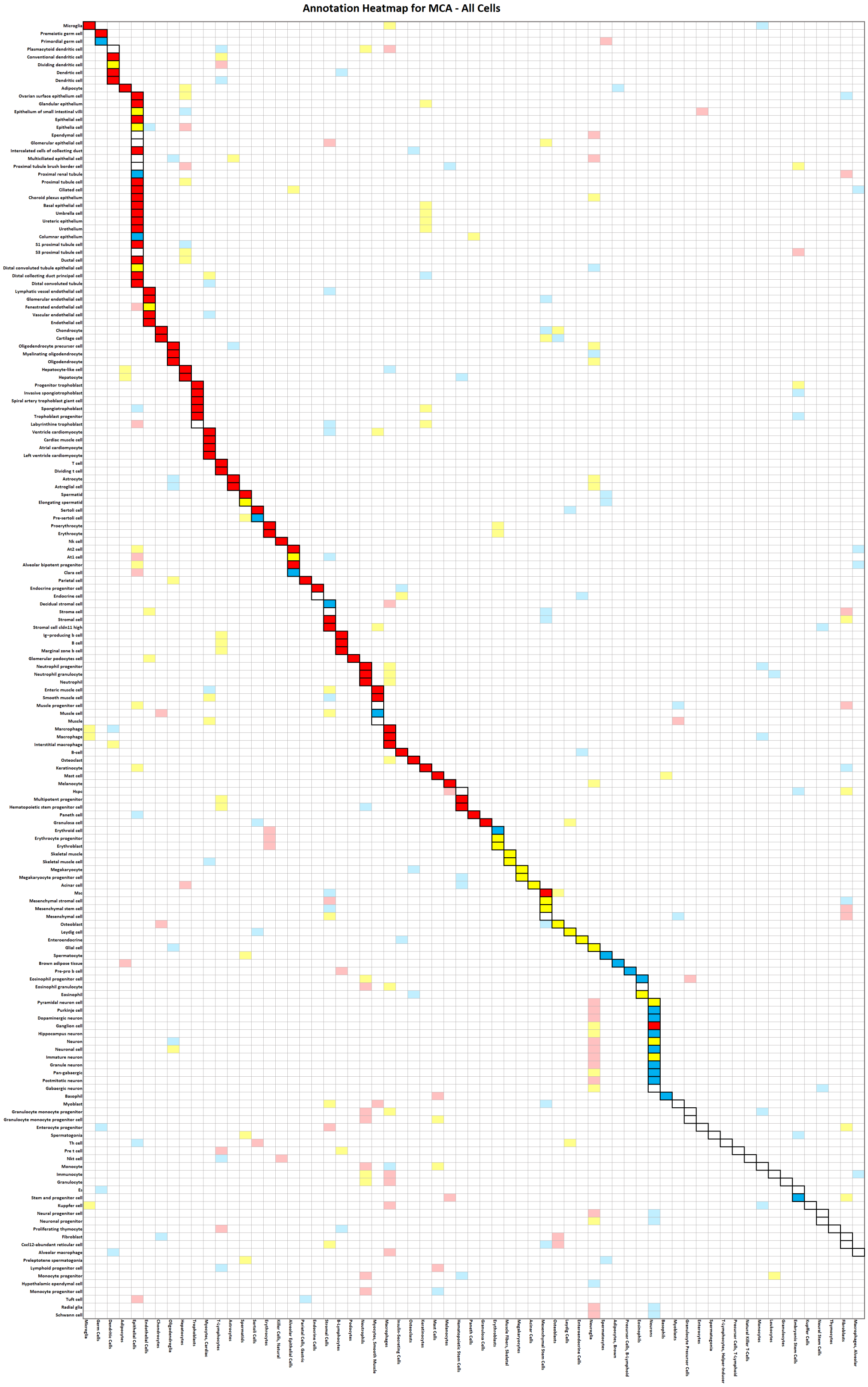

Figure 8: **Annotation Heatmap on MCA Dataset.** The annotation heatmap has its format similar to Fig 6.

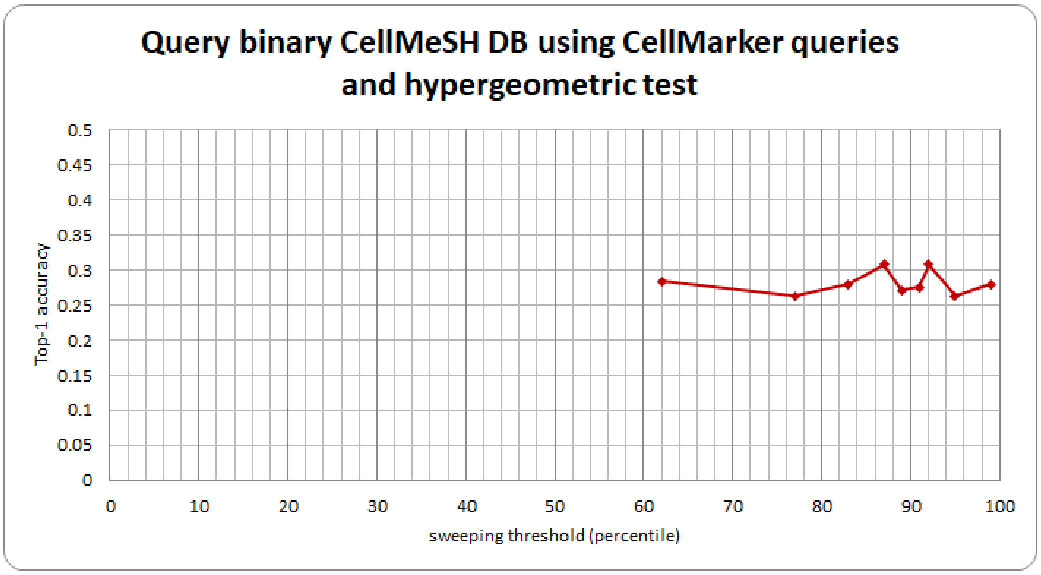


Figure 9: **Selection of Threshold to Binarize CellMeSH Database for Hypergeometric Test.** The hypergeometric test [1] is designed to query binary database, therefore we need to find a proper threshold to binarize the CellMeSH database so that it can be queried by hypergeometric test. We have tried various thresholds. For each threshold, we first get a binary CellMeSH DB by setting its weights above the threshold as 1 and below the threshold as 0. We next prepare 335 queries using CellMarker [2] database so that each query is a set of marker genes for a CellMarker record. We then use the prepared 335 queries to query this binary DB by hypergeometric test, and finally calculate the top-1 accuracy (y-axis). As we can see, the best performance is achieved by the threshold of 87 percentile (87% of the count weights, which are less than 4 in the DB, are set as zero).

(a) Comparison of Different Normalization Methods

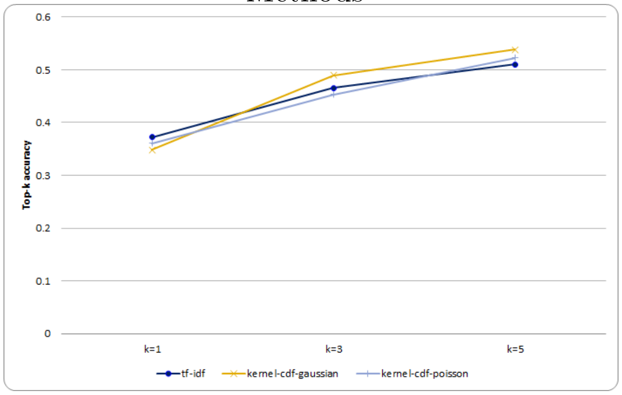


(b) Comparison of flipped and original GSVA

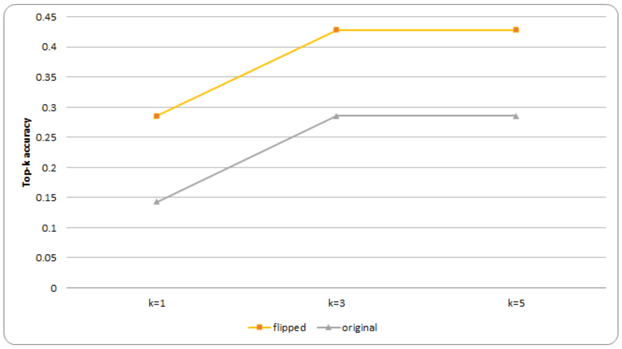


Figure 10: **Settings for GSVA.** (a) The GSVA query method takes input of queries prepared from 335 CellMarker records, and input of normalized CellMeSH database, and predicts cell type for each query. The enrichment score between a candidate cell and the query is a KS-like score. By walking along the sorted genes of the candidate cell, a score gets increased if the gene in the candidate cell overlaps with the query, or gets decreased otherwise. The max score deviated from zero is the enrichment score. The TF-IDF normalization method works better for top-1 accuracy than the kernel-cdf-gaussian and kernel-cdf-poisson normalization methods proposed in GSVA [3]. (b) The GSVA query method takes input of queries prepared from 7 annotated clusters of Zeisel dataset [4], and input of TF-IDF normalized CellMeSH database, and predicts cell type for each query. For the flipped GSVA, the KS-like score between a candidate cell and the query is calculated as follows. By walking along the sorted genes of the candidate cell, a score gets increased if the gene in the candidate cell overlaps with the query (top 100 cluster differentially expressed genes), or gets decreased otherwise. The max score deviated from zero is the enrichment score. For the original GSVA [3], the CellMeSH database is binarized by cutting off 99% weights as zero. The KS-like score between a candidate cell and the query is calculated as follows. By walking along the sorted genes of the query (cluster gene expression), a score gets increased if the gene in the query overlaps with the candidate cell, or gets decreased otherwise. The max score deviated from zero is the enrichment score. We find the flipped mode suits our setting and performs better, so we use this in our main paper evaluation.

[2] Xinxin Zhang, Yujia Lan, Jinyuan Xu, Fei Quan, Erjie Zhao, Chunyu Deng, Tao Luo, Liwen Xu, Gaoming Liao, Min Yan, Yanyan Ping, Feng Li, Aiai Shi, Jing Bai, Tingting Zhao, Xia Li, and Yun Xiao. CellMarker: a manually curated resource of cell markers in human and mouse. *Nucleic Acids Research*, 2018.

[3] Sonja Hänzelmann, Robert Castelo, and Justin Guinney. GSVA: gene set variation analysis for microarray and RNA-seq data. *BMC Bioinformatics*, 14(1):7, 2013.

[4] Amit Zeisel, Ana B. Muñoz-Manchado, Simone Codeluppi, Peter Lönnerberg, Gioele La Manno, Anna Juréus, Sueli Marques, Hermany Munguba, Liqun He, Christer Betsholtz, Charlotte Rolny, Gonçalo Castelo-Branco, Jens Hjerling-Leffler, and Sten Linnarsson. Cell types in the mouse cortex and hippocampus revealed by single-cell RNA-seq. *Science*, 347(6226):1138–1142, March 2015.
